# Evolution of gene regulatory networks by means of selection and random genetic drift

**DOI:** 10.1101/449645

**Authors:** Antonios Kioukis, Pavlos Pavlidis

## Abstract

The evolution of a population by means of genetic drift and natural selection operating on a gene regulatory network (GRN) of an individual has not been scrutinized in depth. Thus, the relative importance of various evolutionary forces and processes on shaping genetic variability in GRNs is understudied. Furthermore, it is not known if existing tools that identify recent and strong positive selection from genomic sequences, in simple models of evolution, can detect recent positive selection when it operates on GRNs. Here, we propose a simulation framework, called EvoNET, that simulates forward-in-time the evolution of GRNs in a population. Since the population size is finite, random genetic drift is explicitly applied. The fitness of a mutation is not constant, but we evaluate the fitness of each individual by measuring its genetic distance from an optimal genotype. Mutations and recombination may take place from generation to generation, modifying the genotypic composition of the population. Each individual goes through a maturation period, where its GRN reaches equilibrium. At the next step, individuals compete to produce the next generation. As time progresses, the beneficial genotypes push the population higher in the fitness landscape. We examine properties of the GRN evolution such as robustness against the deleterious effect of mutations and the role of genetic drift. We confirm classical results from Andreas Wagner’s work that GRNs show robustness against mutations and we provide new results regarding the interplay between random genetic drift and natural selection.

## I. BACKGROUND

### A. Introduction

The path from genotype to phenotype is characterized by an immense number of direct and indirect gene interactions. The relationship between genotype and phenotype has long been of interest to geneticists, developmental biologists and evolutionary biologists. This is partially because the relationship between genotypes and phenotypes is ambiguous and non-linearities appear often. The same phenotype can be produced by a range of genotypes and a single genotype can result in different phenotypes due to the environmental effects [18]. Natural selection operates on various levels of genomic organization, from single nucleotides, genes, networks of genes to complex phenotypes. Phenotypic variation is the first of the three principles required for the action of natural selection [8]. Thus, it may seem inconsistent that tests for the localization of natural selection, *i.e.* selective sweeps, use solely genotypic information, in models that incorporate no gene interactions or genotypic-phenotypic relations. In contrast, they utilize the concept of constant selection coefficient, which can be understood as a summary of the dynamics of the allele under selection, but lacks a clear biological meaning [3]. If a genomic region is identified as the target of positive selection, the next step usually comprises an extensive literature search in an effort to connect the genotype to phenotype, and thus build plausible narratives that explain the action of positive selection [13].

Chevin et al. [3] extended the theory of selective sweeps to the context of a locus that affects a quantitative trait, thus a phenotype, that harbors background genetic variation due to other, unlinked and no-interacting, loci. They assumed a large number of background loci with a small effect on the phenotype. Even though the increase in frequency of a beneficial mutation is slower than the classical one-locus selective sweep, they showed that under such a model, selective sweeps can still be detected at the focal locus, especially if the genetic variation of the background is not too large. Pavlidis et al. [14] showed that when the train under selection is controlled by only a few loci (up to 8 in their simulations), it is possible that an equilibrium is reached, and thus no fixation of an allele. Such an equilibrium scenario occurs more frequently when loci have a similar effect on the phenotype. Contrariwise, if the population is far from the optimum and the focal allele has relatively large effect, then it will reach fixation. In general, multi-locus models allow competition between loci, thus it depends crucially on the initial conditions whether a locus will reach fixation fast, and thus a selective sweep will be produced.

To our knowledge, the first attempt to understand the evolution of regulatory networks was done in the seminal work by Wagner [21]. Wagner evolved numerically a network of genes that assume binary states (either on or off). He studied whether a population of such networks can buffer the (detrimental) effect of mutations after it evolves to reach its optimum. Indeed, he found (Figure 2 in [21]) that after evolving a network of genes by means of natural selection (stabilizing selection), the effect of mutations is considerably lower than a system where evolution has not occurred yet. Natural selection, combined with neutral processes, modifies gene expression and in consequence the properties of GRNs. Ofria et al. [12], using computer simulations, demonstrated that when the mutation rate is greater than zero, selection favors GRN variants that have similar phenotypes. Wagner [22] showed that neutral variants with no effect on the phenotype facilitate evolutionary innovation because they allow for thorough exploration of the genotype space. These ideas can be directly applied to GRNs by employing the concepts of robustness and redundancy. Robustness refers to the resilience that GRNs exhibit with respect to mutations. One mechanism for maintaining robustness is redundancy. Redundancy may be caused by/implemented by gene duplication or by unrelated genes that perform similar functions [11].

**FIG. 1.**
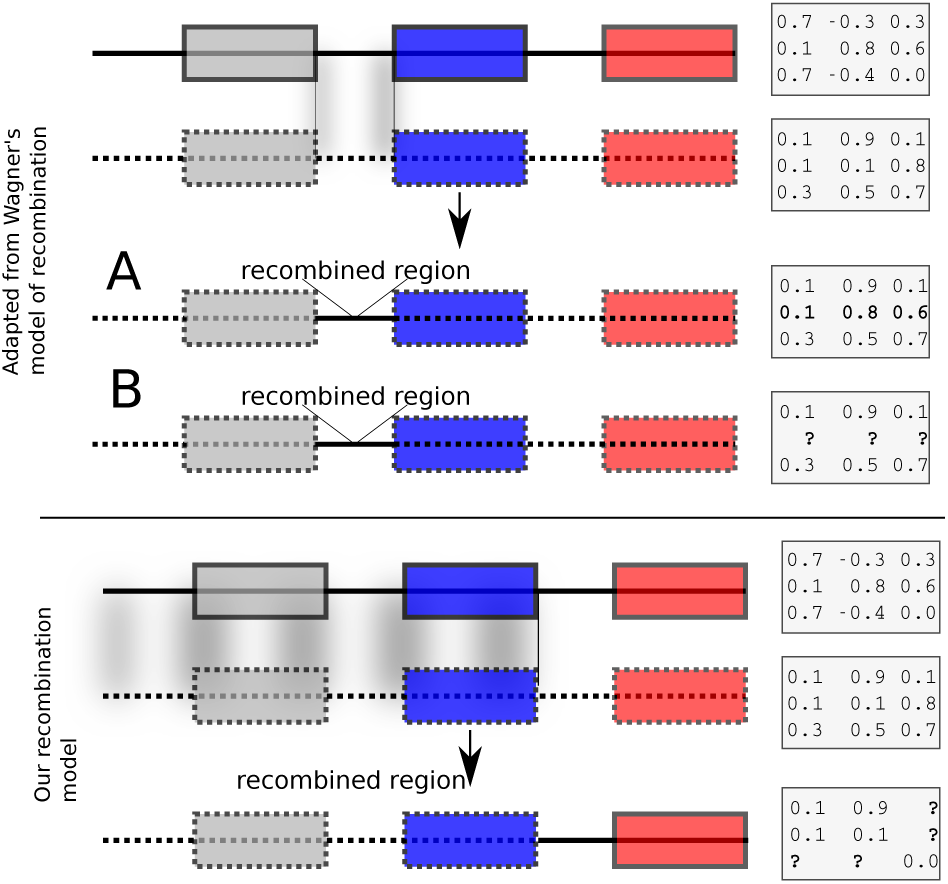
Recombination models implemented by EvoNET. Shaded areas show the gnomic regions that are exchanged due to the recombination process. At the upper panel, Wagner’s model is illustrated, where *cis* regulatory regions can be swapped between individuals of the population. At the bottom panel, our model is shown. In our model, recombination is implemented via a recombination break-point. All genes at its left side inherit both the *cis* and the *trans* regions from one parent, whereas the genes on the right inherit *cis* and *trans* regions from the other parent. The interaction matrix is re-evaluated after recombination.

**FIG. 2.**
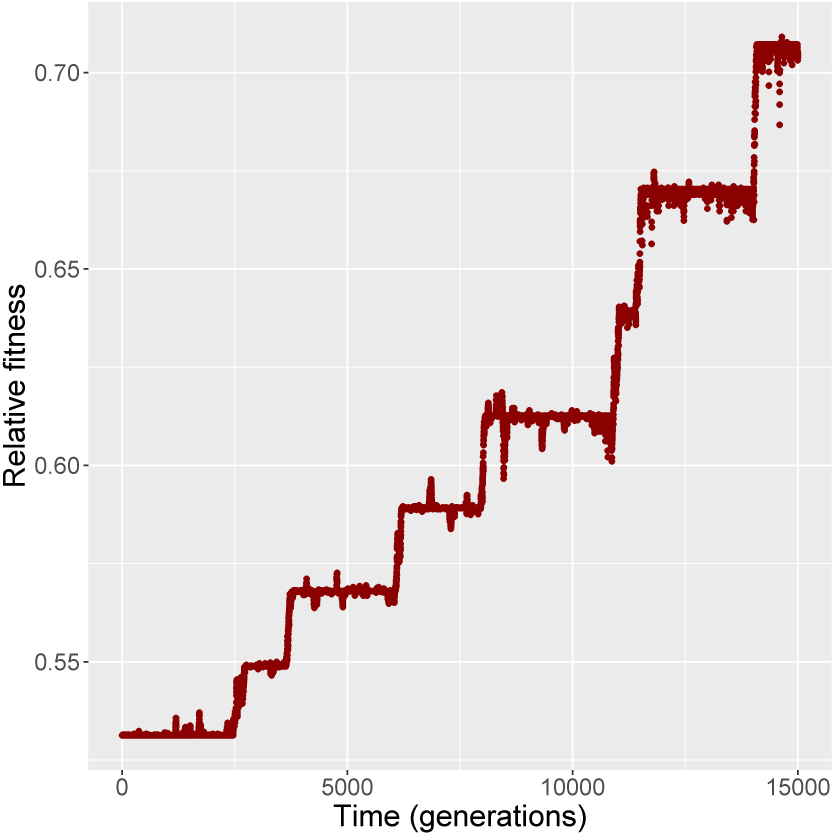
The increment in relative fitness of the population is taking place in discrete steps, in a ladder-like fashion.

Computational tools for detecting positive selection have been developed [1, 10, 15] based on the “hitchhiking” or “selective sweep” theory [9, 20]. Three deviations from classic selective sweep theory are possible because of positive selection effects on GRNs: i) variation in selection intensity through time; ii) soft sweeps that start with several favorable alleles; and iii) overlapping sweeps [4]. Since more than one network configuration can give rise to the same phenotype, the polymorphic patterns at the genome level are not necessarily expected to match the expected polymorphic pattern distribution that is caused by a strong beneficial mutation in just a single, independent gene. This has been shown for selective sweeps on a quantitative trait locus [14]. Adaptation may often be based on pre-existing genetic variation of the population (standing genetic variation), rather than single, new mutations. Thus, it is expected that the new allele may originated from multiple initial alleles, which will in turn weaken the signal of positive selection [17]. Finally, if hitchhiking, as is widely believed, dominates the pattern of neutral diversity, the genome may be subject to multiple overlapping sweeps. Barton [2] has extended earlier branching-process methods to determine how overlapping sweeps reduce mean coalescence time as well as how they reduce the fixation probability of favorable alleles.

In this work, we study via a forward-in-time simulator, named EvoNET, the evolution of a population of GRNs by means of random genetic drift and selection. We extend Wagner’s classical model [21] and subsequent extensions (e.g. [19]) by allowing cyclic equilibria during the maturation period and a different recombination model. We provide results about the robustness of the network to mutations, and its properties during evolution in a fitness landscape (e.g. genetic diversity).

## METHODS

### The model

#### a. Regulatory regions define interactions

We assume a population of *N* individuals. Each individual comprises a set of *n* genes consisting of *cis* and *trans* binary regulatory regions, each of length *L*. A *cis* regulatory region is defined as the region upstream the gene on which other genes of the GRN can bind. Let *R*_*i,c*_ be the *cis* region of the gene *i* and *R*_*j,t*_ the *trans* region of gene *j*. Then, we define a function *I*(*R*_*i,c*_, *R*_*j,t*_) that receives as arguments two binary vectors and returns a float number in the [−1, 1] representing the interaction strength. Negative values model suppression, positive values activation, whereas 0 means no interaction. Any function that takes as arguments binary vectors and returns a value in the [− 1, 1] could be used as the *I* function. Here, for the absolute value of interaction, we use the Equation 1.1:

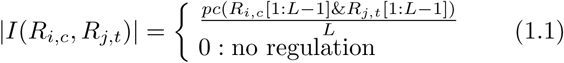

where *pc* is the popcount function, which counts the number of set bits (i.e. 1’s) that are common in the two vectors. The occurrence of interaction, as well as, the + or − *sign*, is defined by the last bit of the *R*_*i,c*_ and *R*_*j,t*_ vectors as:

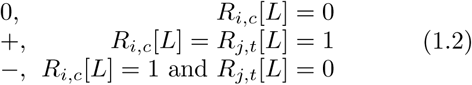

In other words, the first *L*− 1 bits define the strength of the interaction, which is proportional to the number of common set bits (i.e. common 1’s). The last (*L*^*th*^) bit in each vector determines if the interaction is present and if it is suppression or activation. If the last bit of the *cis* element is ‘0’ then it does not ‘accept’ any regulation. If it is ‘1’, then regulation can be either positive or negative, depending on the last bit of the *trans* element.

The above representation of regulation enables a more realistic representation of regulation than Wagner’s model [21] and its more recent extensions [5, 19]. A single mutation in the *cis* region of a gene can affect its regulation by all other genes, and a mutation in the *trans* region of a gene can affect the way it regulates all other genes (see also the section ‘Mutation model of regulatory regions’).

#### b. Interaction matrix and expression levels

Interaction values are stored in a square *M*_*n×n*_ matrix of real values in the [−1, 1], where *n* is the number of genes in the network. A positive *M*_*ij*_ value indicates that gene *j* activates gene *i*, a negative value indicates suppression and 0 represents no interaction. Thus, the row *M*_*i.*_ represents the interaction between all trans regulatory elements and the *cis*-regulatory region of gene *i*. Gene expressions are represented by a vector *E*_*n*_ of *n* elements. In the general case, the expression level *E*_*j*_ of the *j*_*th*_ gene can be a real positive number. Here, however, *E* is a binary vector, indicating only if a gene is switched on or off. Such a representation is more efficient computationally. A similar approach has been used by Wagner [21] and Siegal and Bergman [19].

#### c. Inheritance of regulation and recombination

Each child inherits from its parents (the model allows for two parents or a single mother) the *cis* and *trans* regulatory regions. The initial values of expression levels (at birth) are defined solely by the environment, and here they are initialized to a constant binary vector. If the model allows for two parents, then recombination is possible to occur. We have implemented two recombination models. The first is similar to Wagner [21]’s model that swaps rows of the interaction matrix of parents to form children. Such a model results effectively in exchanging *cis* regulatory elements. Wagner’s model of recombination may be, however, unrealistic because it allows *cis* regulatory regions to be exchanged, however the *trans* regulation (which is modeled by the gene itself) does not change. Thus, the *cis* regions can be exchanged but not the genes associated with the *cis* regions (Figure 1 top panel). In Wagner [21], the interaction values between genes in the recipient and donor genomes remain unchanged after recombination (Figure 1, upper panel A). We implemented Wagner’s model of recombination, but we re-estimated the interaction values between genes in the donor and the recipient genomes. This is necessary because *cis* and *trans* interactions are modified after recombination (Figure 1, upper panel B). We implemented an additional recombination model that allows cross-over events between parental genomes as follows: Assuming that *n* genes exist in the genome (members of the GRN), choose *j*, 0 *< j < n* an integer breakpoint. Then, the first *j* genes inherit both the *cis* and the *trans* regions from one parent, and the last *n*− *j* genes inherit *cis* and *trans* regions from the other parent. The regulation between the first *j* and the last *n*− *j* genes is re-computed from their regulatory regions (Figure 1, bottom panel).

#### d. Mutations

Mutations take place in the *cis* and *trans* regulatory regions during offspring generation. Since regulatory regions are implemented as binary vectors, a mutation can change a position in a region by modifying a 0 to 1 and *vice versa*. On one hand, if a mutation will affect a *cis* region, then all interactions between this *cis* and all *trans* regions might be modified (i.e., the row of the interaction matrix will be affected). On the other hand, if a mutation will change a *trans* region, all interactions between this *trans* and all other *cis* regions might be modified (i.e., the column of the interaction matrix). For each individual, the number of mutations is drawn from a Poisson distribution with parameter *µ* (mutation rate per genome per generation), and then mutations (if any) are placed uniformly among the *cis* and *trans* regulatory regions.

For example, let *R*_*i,cis*_ be the *cis* regulatory region of gene *i* that is going to be mutated. *R*_*i,cis*_ comprises two parts: the [1 : *L* − 1] part, which controls the strength of interactions and the *L* position that controls the type of interaction as described in *Regulatory regions define interactions*. Since mutations in the *L* position may have a dramatic effect, changing the type of interaction (e.g. a repressor might become activator or regulation can be silenced), we implemented two different mutation rates for these two parts of the regulatory regions. Mutations in the first [1 : *L*− 1] part are distributed uniformly. We model with 1% chance the probability that a mutation occurs and the trans region changes its behavior. This modeled the biological fact that mutations that change the nature of an established relationship of two genes is very rare in contrast to changing the strength of the respective relationship.

#### e. Selection

Selection operates on the expression profile of genes. In every generation, selection is applied to select the parents of an individual. Let *E*_*opt*_ represent the optimal vector of expression values for the *n* genes, that is the optimal expression level for the first gene is *E*_*opt*,1_, for the second gene *E*_*opt*,2_ and so on. The fitness of an individual with expression values defined by the *E*_*n*_ vector is defined by:

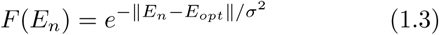

where ‖*E*_*n*_ − *E*_*opt*_ ‖ is a norm of the difference between *E*_*n*_ and *E*_*opt*_ expression vectors (here the Euclidean distance is used). Parents are chosen proportionally to their fitness value *F* (*E*_*n*_).

#### f. Maturation and equilibria

Every ‘new-born individual’ has inherited the regulatory regions from its parents (potentially with mutations) and in addition it has acquired an initial expression vector (expression values for all genes) that is constant for all individuals. Since genes may interact with each other, we have implemented an additional ‘maturation’ process. During the maturation process the expression levels of genes change as a result of gene-gene interactions until either an equilibrium point or a cyclic equilibrium is reached. At the *t* + 1 step of the process a new expression vector *E*_*n*_(*t* + 1) is obtained using the expression vector of the *t*_*th*_ step and the interaction matrix *M* :

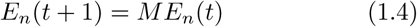

Equivalently, the *i*^*th*^ element 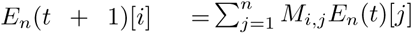. Depending on the interaction matrix *M* and the initial value of the expression vector *E*_*n*_, there are 3 possible outcomes of this process.

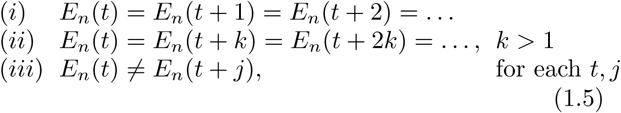

In Wagner’s model [21] as well as in Huerta-Sanchez and Durrett [5], only case (*i*) in Equation 1.5 is considered viable. Case (*i*) facilitates fitness evaluation of the individual using Equation 1.3. Individuals with a maturation process that concludes in (*ii*) or (*iii*) were removed from the population. Here, motivated by Pinho et al. [16] who suggested that in Wagner’s model most networks are cycling, we developed a circadian framework to evaluate the fitness of individuals that conclude in cyclic equilibria during the maturation step. Individuals that conclude in case (*iii*), or individuals that conclude in case (*ii*) but the period *k* is greater than an upper threshold (here 10,000 steps) were considered non-viable and were removed from the population. Thus, if the maturation process concludes in case (*ii*), with *E*_*n*_(*t*) = *E*_*n*_(*t* + *k*) = *E*_*n*_(*t* + 2*k*) = … and *k* > 1, we evaluated the fitness of the individual as the minimum fitness value during the period of a cycle.

## RESULTS

### Comparisons between Neutral Evolution and Selection Scenarios

#### Simulations setup

To explore the gene expression differences between neutral evolution and evolution under directional selection, we simulated neutral datasets and datasets with selection. For the two scenarios, command line arguments were identical except the random number generator seed and the binary flag that denotes whether simulation is neutral. All command lines are provided in the Supplement. Both models were evolved for 15,000 generations. Each individual network comprises 10 genes, each with 30-bit long *cis* and *trans* regulatory elements. The last bit of each regulatory element is responsible for the type of regulation (positive or negative; see Methods) and the remaining 29 bits determine the strength of the interaction, if any. In generation 0, all *cis*-regulatory elements were set to 000 … 01000, that is, initially they can not accept any regulation. In contrast, all *trans*-elements were set to 000 … 01001,*i.e.*, they are activators, thus they can regulate a *cis* element positively (provided that the last bit of the *cis*-element is 1). After maturation (see Methods), the expression vector was converted to binary format (the expression value is 1 if the expression is positive and 0 otherwise). Thus, initially all expression vectors *v* were equal to **0**. The fitness of each person was evaluated after maturation. The optimum was set to the state were all genes were expressed (*i.e.*, state 1 for all genes). For the simulations with selection, the selection intensity 1*/σ*^2^ (see Methods) was set to 1/5. The population size was set to 100 haploid individuals and remained constant throughout the entire simulation.

#### Optimum is gradually reached in a ladder-like fashion

We evaluated whether, and how, the population reaches the optimum state. Given that the initial state was 00000000 (i.e., all genes inactive) and the optimum state was 11111111 (*i.e.*, all genes active), the population had to experience the appropriate changes in its *cis*- and *trans*-regulatory elements, and consequently the GRN, to achieve the activation of all genes. We observed a ladder-like behavior for the average fitness (Figure 2); that is, networks were successively replaced by fitter networks in discrete steps.

At every step of the ‘ladder’, the average population fitness remains approximately constant. After reaching each fitness step, the population starts exploring different GRN topologies until a fitter genotype establishes in the population. While exploring candidate topologies, genetic drift acts and it is therefore possible that the population will not incorporate every novel beneficial network topology that it will encounter. If a beneficial topology overcomes drift, its frequency increases and the average population follows. Finally, when the new topology reaches fixation, the population has reached the next step in the fitness ‘ladder’ (Figure 3).

**FIG. 3.**
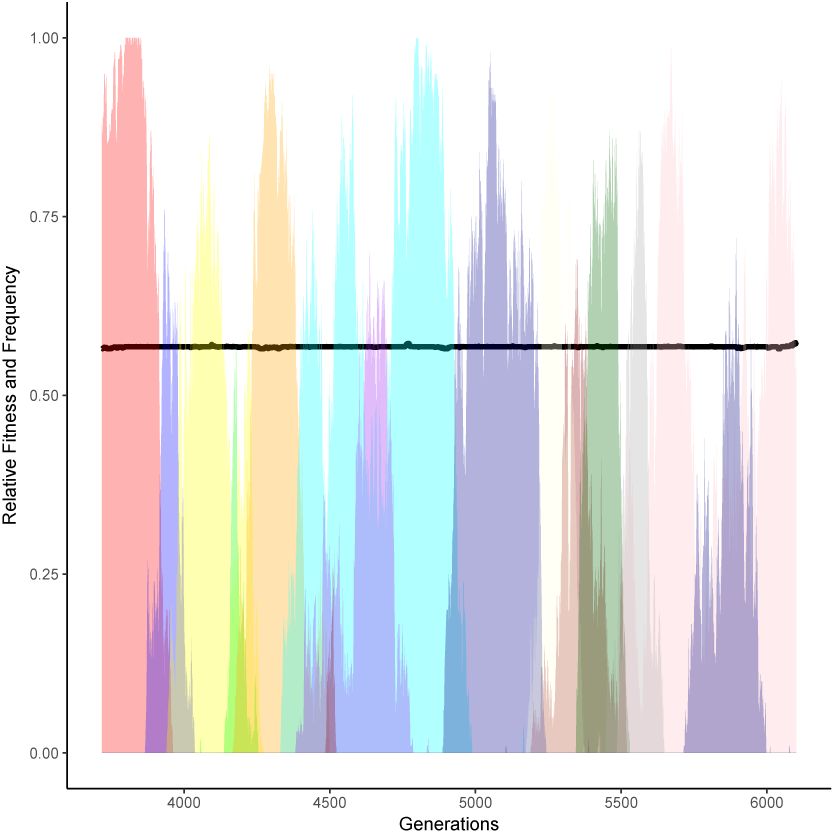
Alternating frequency-trajectories of the various regulatory networks at a certain fitness level (0.5679; black thick horizontal line). All networks have the same fitness. Here, we show only networks with frequency at least 50%. There are 14 different networks.

Mutations and recombination are the driving force behind the exploration of the topology space, since they may result in a novel network topology. By increasing the mutation rate, the number of novel explored topologies increases and waiting times between each step are decreased. (Figure 4).

**FIG. 4.**
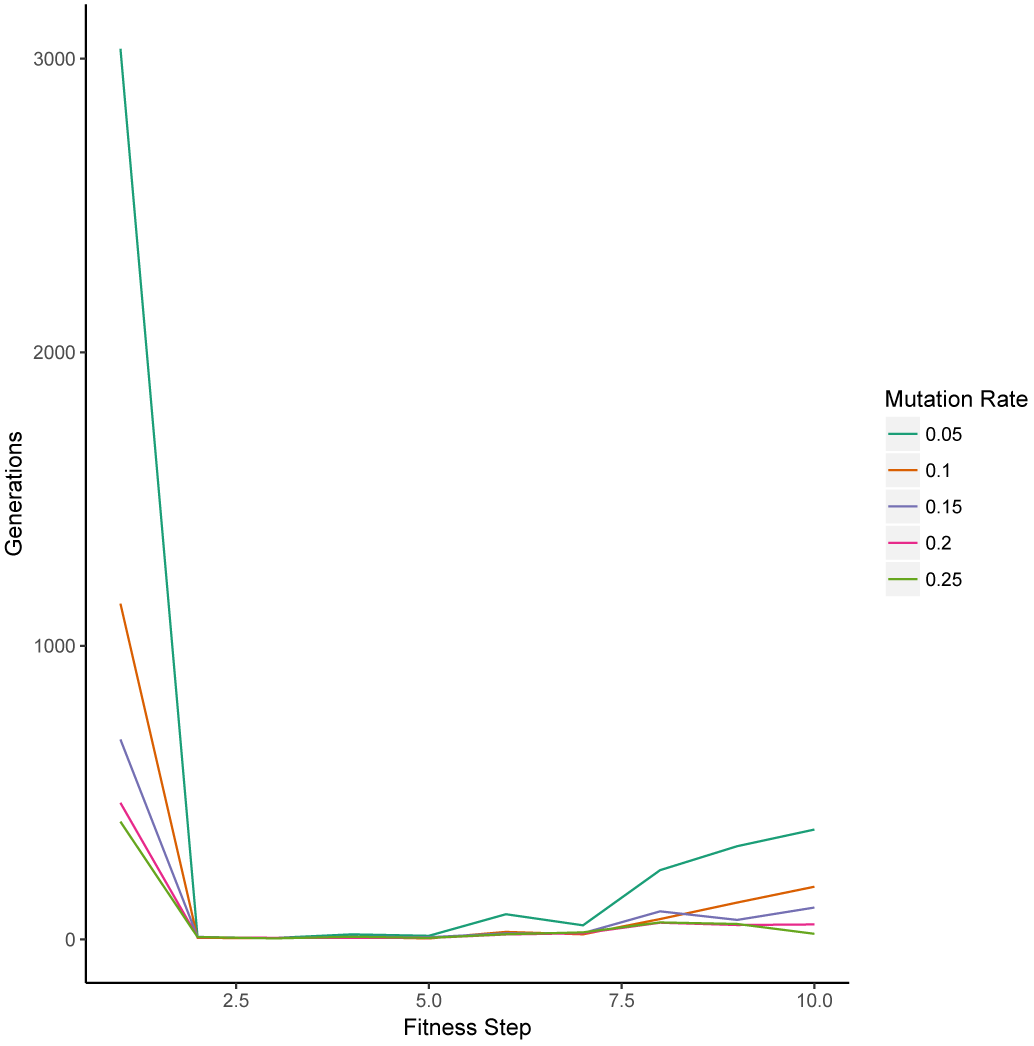
Increasing the mutation rate reduces the time needed to take the next step on the fitness landscape.

Recombination rates also affect the time required for each step. Recombination allows the parental networks to be combined resulting in enhancement of the network variability in the population, thus the optimum can be reached faster. In our simulations our proposed model R1R2 swapping reaches optimum faster than the row-swapping model proposed by Wagner [21] (Figure 5).

**FIG. 5.**
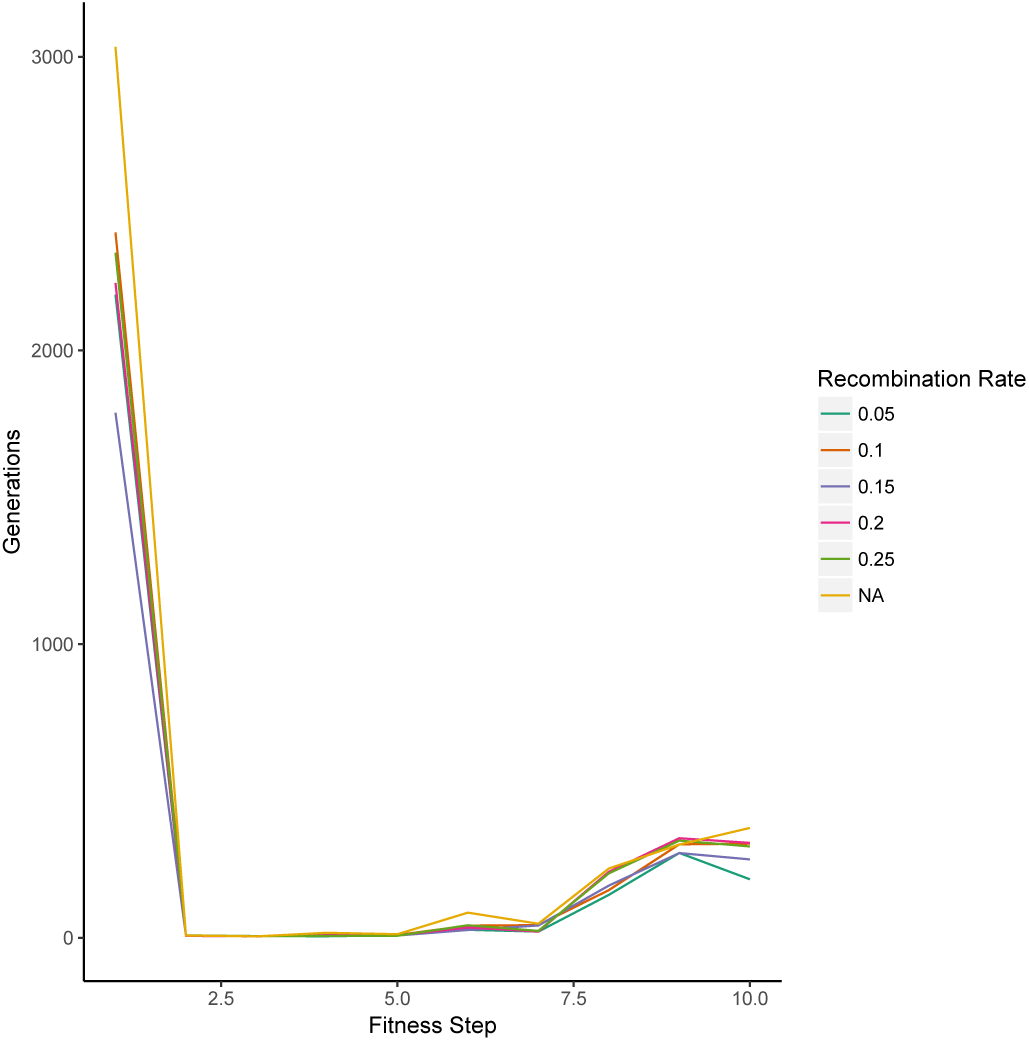
Recombination rates and time needed to take the next step on the fitness landscape. Especially for the first step, which takes most of the time, the least time is achieved when recombination rate is 0.15, i.e. intermediate between the minimum and the maximum.

#### Size of the regulatory space in neutrality and selection

We assessed how the population explores the state space of regulatory networks during its evolution, by evaluating the number of different genotypes individuals obtain. We studied whether neutrality or selection explores the space more efficiently, *i.e.*, which of the two processes allow the population to explore a higher number of genotypes on average.

During the course of evolution, for 15,000 generations, both neutral and selection scenarios experienced a multitude of GRNs. In the selection scenario, the population encountered 17,110 different networks; under neutrality the population experienced only 5,105 GRNs. This means that under selection the population is able to explore a greater part of the space of GRNs than under neutrality. Due to selection pressure, the population moves towards the optimum via genotypes that are optimal at the given time point. Then, due to drift, it explores genotypes with the same fitness (*i.e.*, effectively neutral) until a new optimal genotype overcomes drift and brings the population to the next fitness level.

Under neutrality, the behavior of the population was different. With the selection pressure absent, the fate of genotypes was affected solely by genetic drift. In the limited amount of generations (15,000), and due to the small population size (100 individuals) the population explored a small fraction of the genotypic space centered around the initial state.

### B. Choice of recombination model and shape of fitness landscape affect time to reach optimum fitness

Different optimal states model different fitness landscapes. EvoNET will reach the optimal state regardless of the shape of the fitness landscape. A population following our R1R2 recombination model reaches the optimum faster than a non-recombining population in the cases of the optimal states 1111111111 and 1111100000 (Figure 6). On the other hand, for the optimal states 1100110011 and 1010101010, recombination makes the population reach the optimum slower than the non-recombination scenario.

**FIG. 6.**
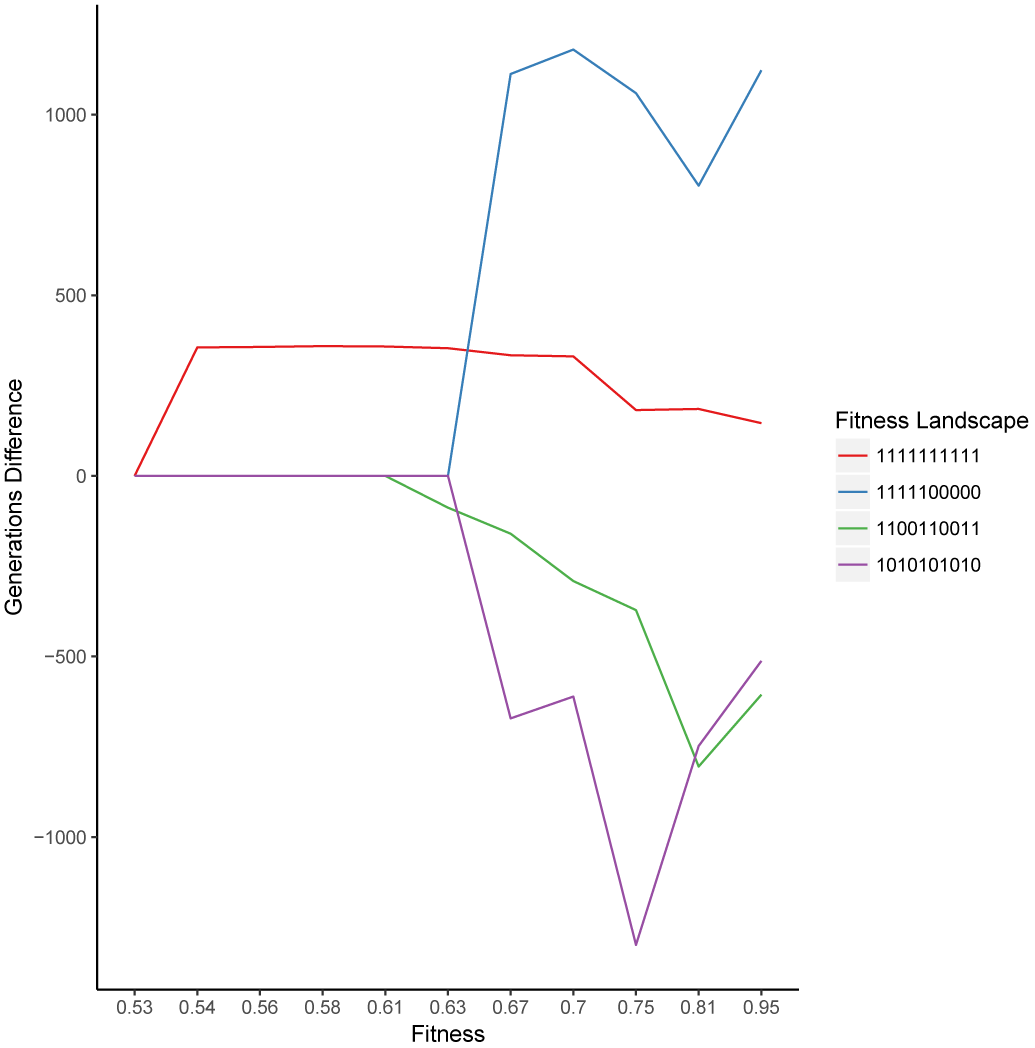
Non-recombining population needs more time to navigate the landscape than recombining population for the 1111111111 and 1111100000 cases. On the other hand the optimum is reached faster for the non-recombining populations when the optimum is set to 1100110011 and 1010101010.

### C. Robustness of Gene Regulatory Network

Robustness to the (phenotypic) effect of mutations has been studied in the framework of GRNs [21], demonstrating that GRNs which reached the phenotypic optimum are less sensitive to mutations, a phenomenon named epigenetic stability. Thus, epigenetic stability was attributed to the evolution of GRNs via the selection process. At discrete time-points EvoNET clones the evolving population (‘core’ population) creating a ‘branch’ population. Each ‘core’ individual has an interactions matrix *M*_*i*_ shared with its ‘clone’. The ‘branch’ population mutates further and then both populations start the maturation progress. The interaction matrices are, then, discretized (*D*_*i*_, *D’*_*i*_).

We assess the GRN robustness at two levels, topology and phenotype. Each GRN has a unique network topology characterizing the strength and effect of all gene interactions. In EvoNET, the topologies are modelled by the interaction matrix, so the additional mutations occurring in the ‘branch’ population have the potential to change the network’s topology. The topology robustness measures whether the ‘core’ and ‘branch’ networks represent the same network topology after the incorporation of the additional mutations on the ‘branch’ population. Phenotypic robustness measures differences in the (binary) expression vector between the two populations after every branching. (Figure 7).

**FIG. 7.**
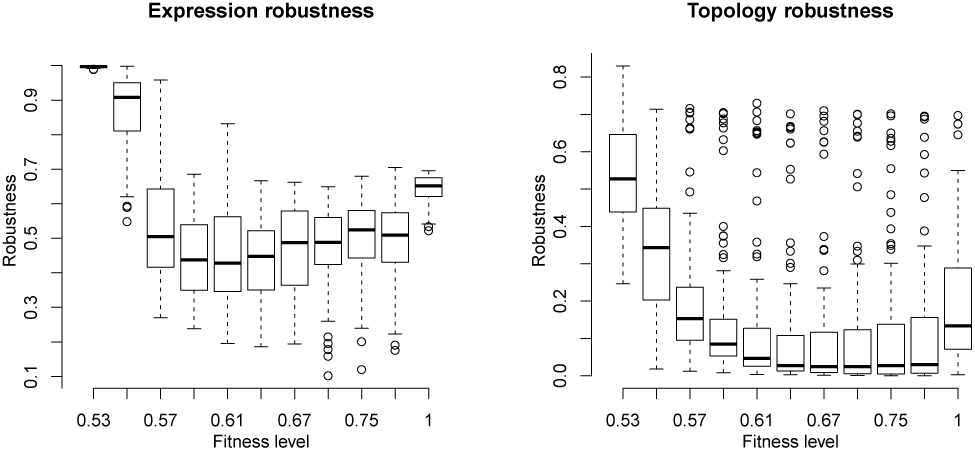
Robustness of the (binary) expression vector and network topology. Initially, the robustness of the expression vector is very high due to the initialization of the simulator. The initial interaction matrix results in the 00 … 0 expression vector. Since no interaction is possible in the beginning, the initial state is robust to mutations. Robustness falls dramatically after the initialization step and increases as fitness increases. The maximum robustness is achieved when the optimum has been reached, on average. The topology is less robust than then expression vector (bottom plot). However, robustness of topology also increases when the population has reached the maximum fitness level.

### D. Effect of neutral genes

All genes in a GRN are not subject to the same evolutionary pressure. Often, a subset of the GRN is evolving under neutrality while other parts are under selection. Using EvoNET we inferred that the number of interactions between neutrally evolving genes and selected genes increases until the population reaches the optimum. When the fitness increases, there are multiple interactions between the two parts (neutral and selected), due to the fact that a beneficial mutation in the neutral part of the GRN has an indirect positive effect on the GRN. In contrast, when the population is at the optimum (right box in 8), mutations are rather deleterious resulting in disadvantageous interactions. Since mutations happen with the same rate across both the neutral and selected part of the GRN, the GRN minimizes the chance that a deleterious mutation will affect it, by discarding the interactions between the different parts (Figure 8).

**FIG. 8.**
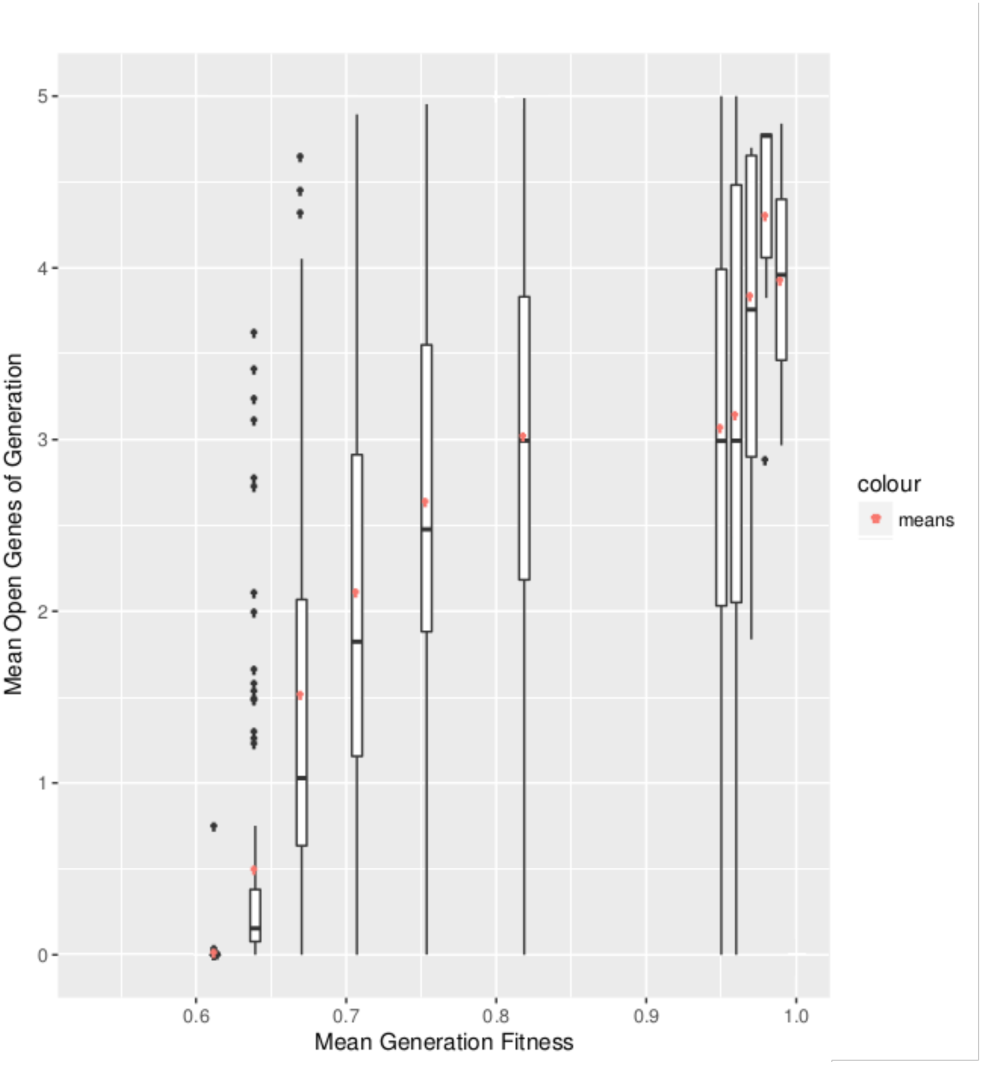
It is beneficial for the GRN to have open connections with neutrally evolving genes outside the GRN when the population is ascending the fitness landscape (boxes; red points represent the means). Upon reaching optimum fitness those interactions tend to be discarded. Boxplots depict the results of 100 simulations, where the majority (blue points) reached each fitness step.

#### 1. Competition Between GRNs of Different Length

We examined whether the size of the GRN is itself a feature on which selection may operate. Thus, we created two distinct GRNs and we let them evolve in the same population. The first GRN, *G*_*s*_, consists of five genes under selection. The second GRN, *G*_*l*_ consists of seven genes. In both GRNs the rest of the genes (five and three, respectively) evolve neutrally. In addition, *G*_*s*_ could not regulate the *trans* region of half of the neutral-evolving genes to simulate a slower mutation rate outside the GRN, whereas the second GRN was free to regulate everything. During the competition between the GRNs, *G*_*l*_ dominated *G*_*s*_ even though *G*_*s*_ had fewer genes under selection so deleterious mutations occurred less frequently. The lack of regulation on the trans-region prohibited *G*_*s*_ from reaching the critical fitness level after which the neutral gene interaction are phased out.

### E. GRN effect

Robustness against mutations is an emergent feature of the GRN [6]. By comparing EvoNET with another algorithm that omits the GRN and directly switches on and off the genes, we demonstrate that the existence of the GRN gives rise to mutational robustness and therefore reaching the fitness optimum faster at high mutation rates. For small mutation rates, robustness and the resulting buffering of mutations happening in EvoNET hinders the acquisition of fitness optimum. When the mutational load increases, however, EvoNET reaches optimum fitness faster due to the robustness created by the GRN. (Figure 9)

**FIG. 9.**
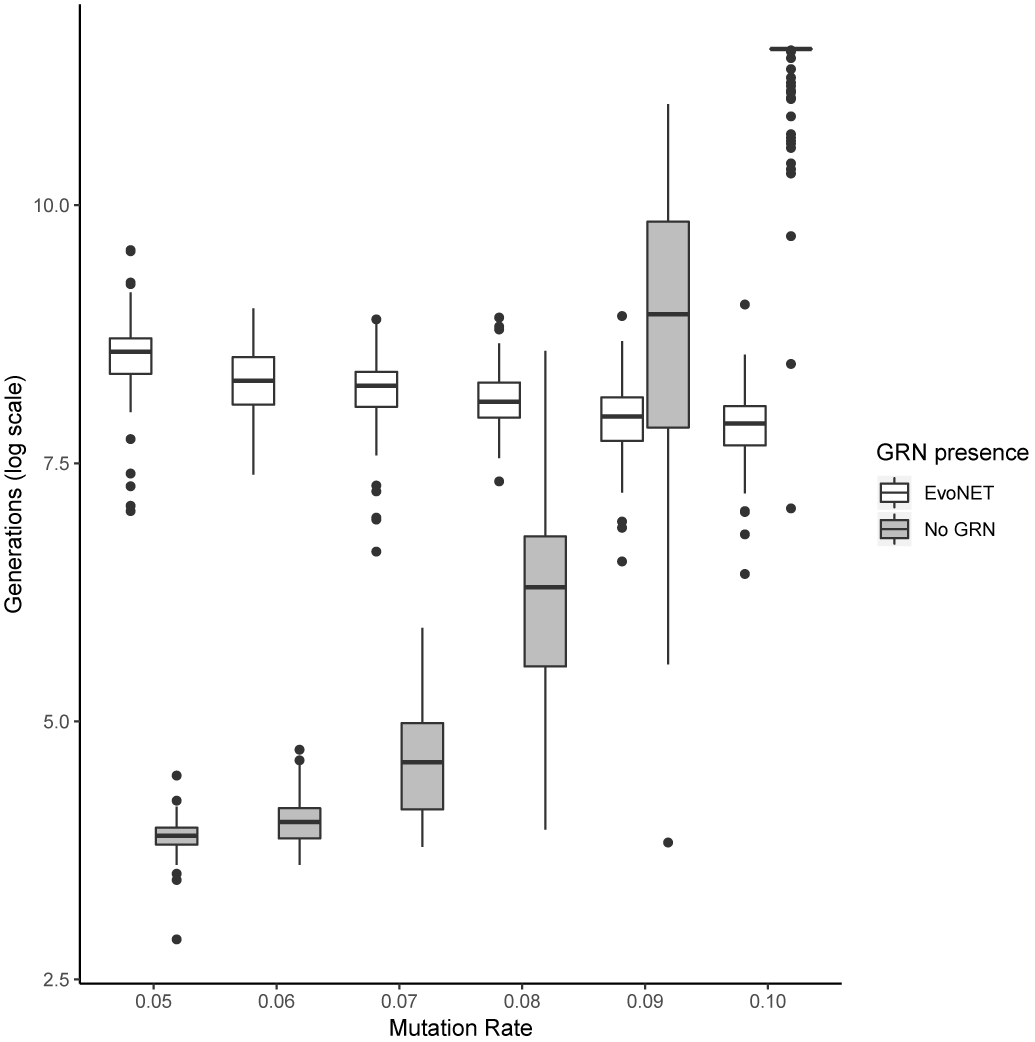
Comparison between the time (in generations) needed to reach the fitness optimum between EvoNET (white) and a similar simulator that directly switches on and/or off genes without employing a GRN (gray boxes). For lower mutation rates, robustness buffers the immediate effect of mutations. Thus, EvoNET reaches the optimum slower than the alternative approach that does not employ GRNs. When the mutation rate increases, mutations, on one hand slow down the simulator without the GRNs. On the other hand, they do not have a detrimental effect on EvoNET due to the buffering effect of the GRN.

## II. DISCUSSION

In this study, we developed EvoNET that creates detailed models of GRNs, thus, enabling the investigation of GRN evolution in the population level. EvoNET extends the algorithm proposed by Wagner [21], by simulating the *cis* and *trans* gene regions creating a more realistic model of the GRN. The regulatory *cis* and *trans* regions interact to create the gene interaction matrix which was the basis of Wagner’s model (Wagner [21] directly mutates the interaction matrix). EvoNET employs the following processes in every discrete generation: birth (with or without recombination), mutation, maturation and fitness calculation. The birth phase is represented by the inheritance of the *cis* and *trans* regions from the previous generation. We introduced a new recombination model (R1R2) that is more realistic than the previously used row-swapping model by [21]. The R1R2 model has a similar behaviour with Wagner’s row swapping model regarding the average time needed for every fitness level (Figure 10). Next, mutations happen, affecting the *cis* and *trans* regions. *cis* and *trans* regions interact to create a new interaction matrix. EvoNET models the type of interaction using the formula shown in Equation 1.1. In the maturation phase, the phenotype is obtained. In contrast to previous studies, we handled the cyclic equilibria instead of discarding them [21] and we evaluated their fitness, making the evolution model more realistic.

**FIG. 10.**
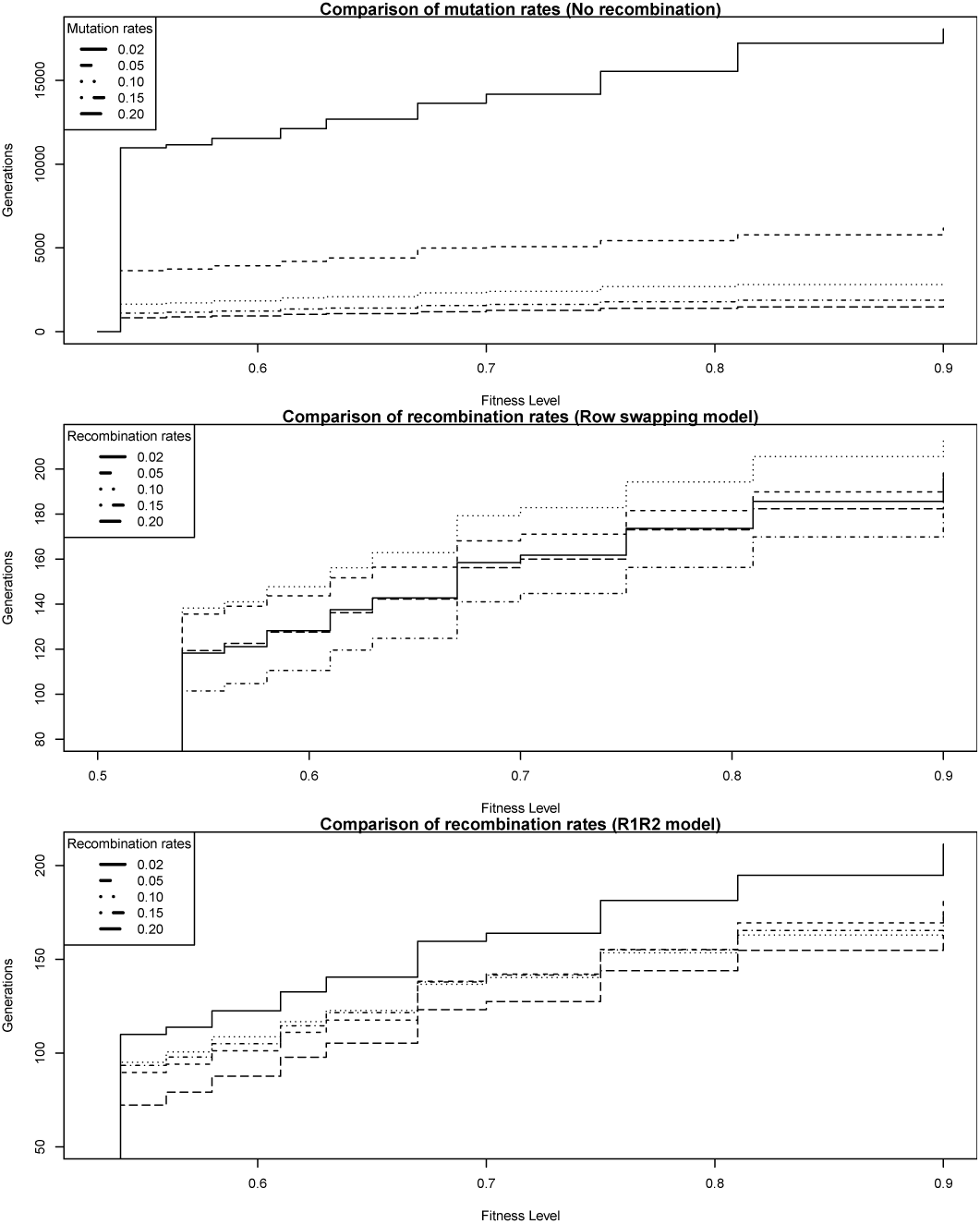
Comparison between the no-recombination model, the Wagner’s swapping and the R1R2 recombination model regarding the generations required to reach optimum fitness.

In the simulations where the mutation rate is sufficiently low, we observed that the fitness landscape takes a ladder-like shape. The steps of the ladder represent the time (measured in generations) that the population explores the genotype space by searching different network topologies allowing for the next step in the fitness ‘ladder”. Small increases in the mutation and recombination rates result in a decrease in the exploration time (Figure 10) due to the increased number of mutations permitting a quicker exploration of candidate network topologies.

We explored the role of robustness of the GRNs while they undergo selection. Robustness is important because it verifies the existence of phenotypically neutral mutations and allows for complex biological structures that are robust to the detrimental effects of mutations. There are two robustness levels acting as canalization attempts, the network topology and the phenotype. Phenotype is more robust to mutations than network topologies, since topology is more directly related to the regions affected by mutations. By comparing EvoNET with a GRN-less simulation program we conclude that these robustness levels permit the GRN to increase its fitness even under high mutation rate. In lower mutation rates, robustness acts as a barrier on the effect of all mutations driving the population to a flat network space thus avoiding perturbations [7]]. In contrast, when the mutation rate increases, the GRN robustness limit is overcome and deleterious mutations, which are more frequent, are immediately affecting the network. GRNs are able to buffer the detrimental effect of mutations, highlighting their biological significance.

Each GRN interacts with other genes which may be evolving under neutrality. By using EvoNET to simulate neutrality and selection acting on parts of the GRN we can draw conclusions on these interactions’ effect. While those interactions are beneficial as the population increases its fitness level, they are discarded from the population when it reaches the fitness optimum. A plausible explanation is that the GRN manages to achieve higher robustness level by removing unnecessary genes and also avoids the effect of deleterious mutations happening on the additional genes of the GRN.

## III. CONCLUSION

Gene Regulatory networks play a vital role in the development of evolutionary advantageous traits for all organisms. In this study we have presented EvoNET, a versatile simulator for the evolution of GRNs through means of genetic drift and selection. Through the use of EvoNET we were able to identify new levels of genetic robustness as well as verify the findings of previous research. The source code of EvoNET and its manual are freely available from https://github.com/antokioukis/evonet.

## References

[1] N. Alachiotis, A. Stamatakis, and P. Pavlidis. OmegaPlus: A scalable tool for rapid detection of selective sweeps in whole-genome datasets. Bioinformatics, 28(17), 2012. ISSN 13674803. doi:10.1093/bioinformatics/bts419.

[2] N. H. Barton. Linkage and the limits to natural selection. Genetics, 140(2):821–841, jun 1995. ISSN 0016-6731. URL http://www.ncbi.nlm.nih.gov/pubmed/7498757.

[3] L.-M. Chevin et al. Selective sweep at a quantitative trait locus in the presence of background genetic variation. Genetics, 180(3):1645–1660, 2008.

[4] J. Hermisson and P. S. Pennings. Soft sweeps: molecular population genetics of adaptation from standing genetic variation. Genetics, 169(4):2335–2352, apr 2005. ISSN 0016-6731. doi:10.1534/genetics.104.036947. URL http://www.genetics.org/cgi/content/abstract/169/4/2335http://www.ncbi.nlm.nih.gov/pubmed/15716498.

[5] E. Huerta-Sanchez and R. Durrett. Wagner’s canalization model. Theoretical population biology, 71 (2):121–30, mar 2007. ISSN 0040-5809. doi:10.1016/j.tpb.2006.10.006. URL http://www.ncbi.nlm.nih.gov/pubmed/17178139.

[6] A. Krishnan, M. Tomita, and A. Giuliani. Evolution of gene regulatory networks: Robustness as an emergent property of evolution. Physica A: Statistical Mechanics and its Applications, 387(8-9):2170–2186, 2008.

[7] R. E. Lenski, J. E. Barrick, and C. Ofria. Balancing robustness and evolvability. PLoS biology, 4(12):e428, 2006.

[8] R. C. Lewontin. The units of selection. Annual review of ecology and systematics, 1(1):1–18, 1970.

[9] J. Maynard Smith and J. Haigh. The hitch-hiking effect of a favourable gene. Genetical research, 23(1):23–35, feb 1974. ISSN 0016-6723. URL http://www.ncbi.nlm.nih.gov/pubmed/4407212.

[10] R. Nielsen, S. Williamson, Y. Kim, M. J. Hubisz, A. G. Clark, and C. Bustamante. Genomic scans for selective sweeps using SNP data. Genome research, 15(11):1566–75, nov 2005. ISSN 1088-9051. doi:10.1101/gr.4252305. URL http://www.pubmedcentral.nih.gov/articlerender.fcgi?artid=1310644{&}tool=pmcentrez{&}rendertype=abstract.

[11] M. A. Nowak, M. C. Boerlijst, J. Cooke, and J. M. Smith. Evolution of genetic redundancy. Nature, 388(6638):167–171, jul 1997. ISSN 0028-0836. doi:10.1038/40618. URL http://www.ncbi.nlm.nih.gov/pubmed/9217155.

[12] C. Ofria, C. Adami, and T. C. Collier. Selective pressures on genomes in molecular evolution. Journal of Theoretical Biology, 222(4):477–483, jun 2003. ISSN 0022-5193. URL http://www.ncbi.nlm.nih.gov/pubmed/12781746.

[13] P. Pavlidis, J. D. Jensen, W. Stephan, and A. Stamatakis. A critical assessment of storytelling: gene ontology categories and the importance of validating genomic scans. Molecular biology and evolution, 29(10):3237–3248, 2012.

[14] P. Pavlidis, D. Metzler, and W. Stephan. Selective sweeps in multilocus models of quantitative traits. Genetics, 192 (1):225–239, 2012.

[15] P. Pavlidis, D. Živković, A. Stamatakis, and N. Alachiotis. SweeD: Likelihood-based detection of selective sweeps in thousands of genomes. Molecular Biology and Evolution, 30(9), 2013. ISSN 07374038. doi: 10.1093/molbev/mst112.

[16] R. Pinho, E. Borenstein, and M. W. Feldman. Most networks in wagner’s model are cycling. PloS one, 7(4): e34285, 2012.

[17] M. Przeworski, G. Coop, and J. D. Wall. The signature of positive selection on standing genetic variation. Evolution; International Journal of Organic Evolution, 59(11):2312–2323, nov 2005. ISSN 0014-3820. URL http://www.ncbi.nlm.nih.gov/pubmed/16396172.

[18] R. Sansom and R. N. Brandon. Integrating evolution and development: From theory to practice. MIT Press, 2007.

[19] M. L. Siegal and A. Bergman. Waddington’s canalization revisited: developmental stability and evolution. Proceedings of the National Academy of Sciences, 99(16):10528–10532, 2002.

[20] W. Stephan, T. H. E. Wiehe, and M. W. Lenz. The effect of strongly selected substitutions on neutral polymorphism:Analytical results based on diffusion theory. Theoretical Population Biology, 41(2): 237–254, apr 1992. ISSN 0040-5809. doi:10.1016/0040-5809(92)90045-U. URL http://www.sciencedirect.com/science/article/B6WXD-4F1Y9N0-3M/2/1245281bba0c6b542457fdd75c343edf.

[21] A. Wagner. Does evolutionary plasticity evolve? Evolution, 50(3):1008–1023, 1996.

[22] A. Wagner. Neutralism and selectionism: a network-based reconciliation. Nature Reviews. Genetics, 9 (12):965–974, dec 2008. ISSN 1471-0064. doi:10.1038/nrg2473. URL http://www.ncbi.nlm.nih.gov/pubmed/18957969.

